# De novo protein backbone generation based on diffusion with structured priors and adversarial training

**DOI:** 10.1101/2022.12.17.520847

**Authors:** Yufeng Liu, Linghui Chen, Haiyan Liu

**Affiliations:** MOE Key Laboratory for Membraneless Organelles and Cellular Dynamics, School of Life Sciences, Division of Life Sciences and Medicine, University of Science and Technology of China, Hefei, Anhui 230027, China; Oristruct Biotech Co., Ltd; Biomedical Sciences and Health Laboratory of Anhui Province, University of Science and Technology of China, Hefei, Anhui 230027, China; School of Data Science, University of Science and Technology of China, Hefei, Anhui 230027, China

## Abstract

In de novo deisgn of protein backbones with deep generative methods, the designability or physical plausibility of the generated backbones needs to be emphasized. Here we report SCUBA-D, a method using denoising diffusion with priors of non-zero means to transform a low quality initial backbone into a high quality backbone. SCUBA-D has been developed by gradually adding new components to a basic denoising diffusion module to improve the physical plausibility of the denoised backbone. It comprises a module that uese one-step denoising to generate prior backbones, followed by a high resolution denoising diffusion module, in which structure diffusion is assisted by the simultaneous diffusion of a language model representation of the amino acid sequence. To ensure high physical plausibility of the denoised output backbone, multiple generative adversarial network (GAN)-style discriminators are used to provide additional losses in training. We have computationally evaluated SCUBA-D by applying structure prediction to amino acid sequences designed on the denoised backbones. The results suggest that SCUBA-D can generate high quality backbones from initial backbones that contain noises of various types or magnitudes, such as initial backbones coarsely sketched to follow certain overall shapes, or initial backbones comprising well-defined functional sites connected by unknown scaffolding regions.

## 1 Introduction

De novo protein design possesses enormous potential in biotechnology and biomedicine. Conventionally, computational protein design is carried out by performing energy-guided optimizations [1, 2, 3, 4]. Despite a number of breakthroughs [5, 6], this approach suffers from accuracy problems (associated with the empirical nature of the energy functions) as well as sampling problems (associated with the intrinsic difficulties of exploring rough and enormous energy landscapes). To circumvent these difficulties, machine learning methods, and more recently deep learning methods, are being increasingly applied [7, 8, 9]. For structure-based sequence design, deep learning methods achieving outstanding native sequence recovery rate have been developed [10, 11, 12, 13, 14, 15], with several methods demonstrated to be much more robust and more accurate in wet experimental tests [16, 17] than conventional energy minimization approaches. The more challenging problem of de novo structure design is also being actively investigated [18, 7, 4, 19]. Among the various methods reported so far, at least two have already been shown (with experimentally solved protein structures) to be able to design proteins of novel, non-natural structures: one is based on repurporting structure prediction methods in a novel iterative sequence updating process called “hallucination” [18, 19]; the other is to use Langevin dynamics to optimize protein backbones described by a neural network-form statistical energy function named SCUBA (for sidechain unknown backbone arrangement) [4].

The principal idea of SCUBA is to use a data-driven model to generate designable backbones at high resolutions prior to selecting amino acid sequences[4]. This idea provides a basic framework of de novo backbone design that is orthogonal to structure prediction models such as AlphaFold2[20] or RosettaFold[21]. However, the backbone-centred total energy of SCUBA is only approximated by a linear combination of separately-trained small neural networks, each of which uses a manually crafted set of structural features of only a small number of residues. This approximation restricts the form of the learned model and may lead to substantial amount of information loss or distortion. Moreover, the Cartesian space Langevin dynamics used by SCUBA is computationally inefficient: the simulations require tens of thousands of steps, and are prone to getting trapped in local minima. In principle, these weaknesses of the original SCUBA protocol can be overcome by leveraging the latest developments in deep learning.

A number of recent studies have explored protein structure generation with a variety of deep learning approaches[7, 9] including the Variational Auto Encoder (VAE) [22, 23], generative adversarial network (GAN) [24, 25, 26] and the denoising diffusion probability model (DDPM)[27, 28]. Most of them adopted the strategy of first generating 2-D images depicting residue-pairwise geometric features, and then using these features as restraints in subsequent energy-guided optimization of the 3-D structure[29]. With this approach, the accuracy and computational efficiency may be ultimately limited by the energy minimization protocol. To avoid this, a more desired approach is to use networks that directly output 3-D structures (e.g., by using the mechanism invented by AlphaFold2 for end-to-end protein structure prediction[20]). DDPMs, the type of generative models that achieved immense recent successes in image and audio generations[30, 31, 32, 33], may by adopted for this task. However, unlike the tasks of image or audio generation, for which the final results are in silico data intended for human interpretations, the tasks of protein design must produce ultimate results that correspond to autonomous, physically realizable entities[34]. Thus the physical plausibility of the generated protein structures must be emphasized.

In the current study, we developed SCUBA-D, a diffusion-based deep learning network for de novo protein structure design. As SCUBA, the objective of SCUBA-D is to transform a low quality (i.e. undesignable) initial input backbone into a high quality (i.e. designable) denoised backbone, which should be of sufficiently high resolution for subsequent amino acid sequence selection. We have developed SCUBA-D by starting from a basic network architecture that implements DDPM of protein structures with the pair-representation updating and invariant point attention (IPA) algorithms of AlphaFold2. Then additional components are gradually introduced into the model to improve the physical plausibility of the denoised backbone. The final network comprises first a module for lower-resolution structure denoising, followed by a structure diffusion module for denoising to higher resolutions, the output of the former module used by the latter as priors. In the higher resolution module, the structure diffusion is assisted by the simultaneous diffusion of a pre-trained language model (LM) representation of the amino acid sequence. Additionally, the training losses of SCUBA-D have been defined by using multiple GAN-style discriminators alongside with usual structure deviation losses. We have computationally evaluated SCUBA-D by applying structure prediction to amino acid sequences designed on the denoised backbones. The results suggest that SCUBA-D can generate high quality backbones from input initial backbones that contain noises of various types or magnitudes, such as initial backbones coarsely sketched to follow certain (predefined) overall shapes, or initial backbones comprising well-defined functional sites connected by unknown scaffolding regions.

## 2 Results

### 2.1 An overview of SCUBA-D

Figure 1 shows the overall framework of our model. More details are given in the Method section. Briefly, the end-to-end model has three main components: a lower-resolution denoising module (LRDM) for generating a prior structure (PS) from an initial structure (IS), a LM-assisted structure diffusion module (SDM) for generating a denoised output structure (OS) at higher resolutions, and discriminator networks providing extra adversarial losses for training the denoising diffusion module.

**Figure 1.**
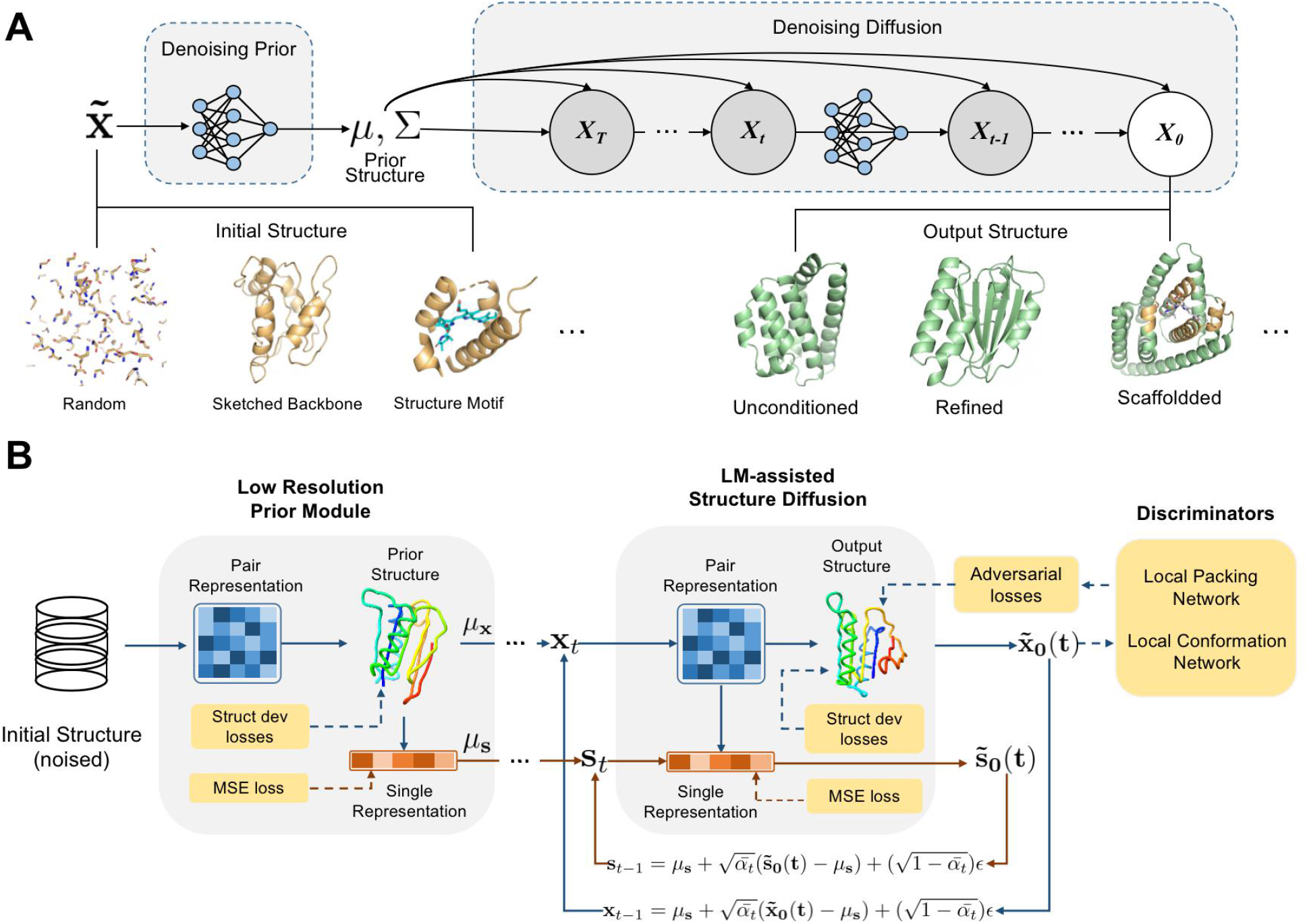
An overview of the SCUBA-D model. **A**, The denoising process using PriorDDPM. Starting from an initial structure, a prior structure is generated by a denoising network. The prior structure is used as the non-zero mean of the starting structure for the reverse deffusion process modelled by DDPM to yield a high quality denoised output structure. Different types of initial structures can be provided so that the output structure can satisfy the various design specifications in different tasks. **B**, The framework of SCUBA-D. The main modules and data representations are illustrated. The arrows show the data flow, with dashed arrows showing the data flow for training only.

In this framework, the initial structures (ISs) can be completely random (for unconditioned structure generation), or it may be utilized to implement various design objectives or constraints on the output or generated structures (see Figure 1A). For example, to design backbones of a pre-specified overall topology, a sketched backbone that coarsely follows a given topology [4] can be used as the IS. Alternatively, an IS can contain well-defined parts which serve as motifs to be integrated into the designed backbones.

To train an LRDM that can process the different types of ISs, during training the ISs have been produced by perturbing natural protein structures in a variety of ways, including randomly masking parts of the structures, adding random noises to the positions and orientations of residues or secondary structure elements, and sampling using SCUBA-driven, high-temperature stochastic simulations[4].

Internally, the LRDM uses a network comprising operations similar to those of the evoformer and invariant point attention (IPA) algorithms of AlphaFold2[20] to update a residue pair-representation of the 3-D structure, and to produce denoised amino acid locations and orientations. The objective of this network is to produce a coarse backbone (the prior structure or PS) that has been improved over the IS, but preserve the topology information of the IS (if the IS contains any). Any well-defined part of the IS (for example, a predefined structural motif) can be marked to be strictly preserved in the PS (Interestingly, if the entire IS is a natural backbone, the LRDM can effectively preserve the entire backbone without using any mark).

The SDM takes the output PS of the LRDM and refines it by using a series of denoising steps, which can be numbered from T to 0, T being an integer denoting the total number of steps. At step t, an internal denoising network is used in the SDM to estimate an output structure for that step (noted as 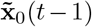) from a noised version of the output structure estimated by the previous step (noted as 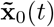) and the PS. This denoising subnetwork has essentially the same structure as the LRDM (but with different numbers of internal nodes and different values of parameters). The noises added to 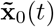 are normally distributed noises with step number-dependent variances.

A language model (LM) of the amino acid sequence (the ESM1b model [35]) is used to assist the structure diffusion process in the following way. In LRDM, a 1-D or single representation (noted as *μ*_*s*_) is predicted from the PS by using a geometric vector perception (GVP) [36] network. This representation (noted as **s**_*t*_) is co-diffused with **x**_*t*_ in SDM. Inside the SDM, the denoising transformation from 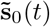 to 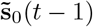 is made to depend on the pair-representation of 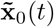.

The LRDM has been trained by minimizing structure deviation losses of the PS with respect to the natural structure together with the mean square errors (MSE) of the predicted sequence representation (*μ*_*s*_) with respect to the LM representation of the native amino acid sequences. The losses for training the SDM comprise structure deviation losses with respect to the natural structure supplemented with adversarial losses (see below) as well as the MSE loss of the LM sequence representations.

The adversarial losses mentioned above are determined by using two discriminator networks. One is a graph-based packing discriminator, which evaluates the plausibility of the local structural environments of individual residues in the OS. The other is a backbone torsion correlation discriminator, which evaluates the plausibility of the backbone conformation represented by contiguous torsional angles. The discriminators have been trained jointly with the LRDM and SDM by using the GAN strategy with feature-matching losses [37].

### 2.2 The effects of different components of the model

We used 25 natural protein domains (see Method) to test our model and to illustrate the effects of the different components of our model. These domains belong to different categories of CATH topology [38], and do not share topology categories with the training structures.

First, we considered the effects of using the LM sequence representation diffusion to assist structure diffusion (without the GAN losses) in the SDM. Four different models are compared. One is the base model, which uses structure diffusion only without considering the LM sequence representation. The other three models predicts three different LM representations (see the caption of Figure 2A) of the native amino acid sequence from the denoised structures. For fair comparisons, the four models have been trained using the same number of steps, at which the training losses were all close to their respective plateau values.

**Figure 2.**
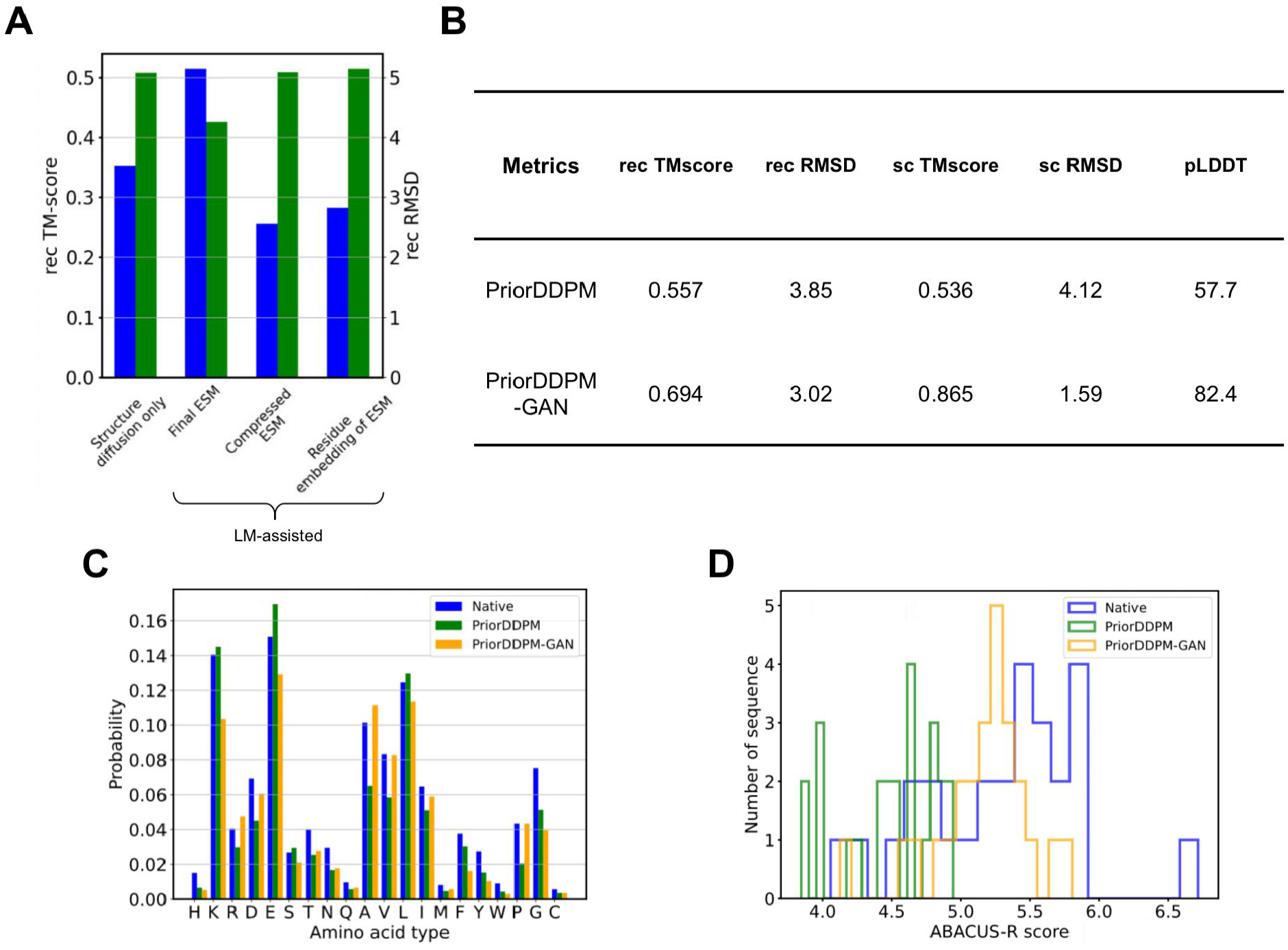
The effects of incorporating sequence language models and GAN losses in SCUBA-D. **A**, The effects of incorporating sequence representations. The results reflect deviations of the output structures from initial structures when natural protein structures are used as input. The averaged RMSDs(green) and TM-scores (blue) from tests using 25 natural protein structures are shown. The four models compared in this figure have been trained with an equal number (10K) of steps. The structure diffusion only model has been trained without incorporating sequence representation. The three LM-assisted models have been trained by using different single representations of the sequence. “Final ESM” used the representation from the last layer of the ESM1b model; “Compressed ESM” used a compressed representation from the final ESM (see Methods); and “Residue embedding of ESM” used the embedding vectors for amino acid types of ESM1b. **B**, The effects of the GAN-style losses. Here “PriorDDPM” and “PriorDDPM+GAN” denote respectively the models trained without and with the GAN losses. Results are from the same tests using 25 natural protein structures as the initial structures. The recRMSDs and recTM-scores are deviations of the output structures from the intial structures, while the scTM-scores and scRMSDs are the deviations of the output structures from the AlphaFold2-predicted structures using ABACUS-R-designed sequences on the output structures. **C**, The amino acid type distributions in ABACUS-R sequences designed on different backbone structures (“Native” stands for the 25 natural structures, while “PriorDDPM” and “PrriorDDPM” stand for output structures of the respective networks). **D**, The distributions of the per-residue logits scores of ABACUS-R sequences designed on different backbone structures.

The 25 natural structures were used as ISs. They were almost unchanged after passing through the LRDM. The deviations of the OSs from the respective ISs generated by the four different models are compared in Figure 2A. The LM-assisted model that predicts the “final ESM” representation substantially reduced the averaged deviations of the OSs from the ISs (here the natural structures) relative to the base model, while the other LM-assisted models exhibit performances similar to or poorer than the base model according to this metric. Thus below we only consider the LM-assisted model with the “final ESM” representation.

Then, we examined the effects of the GAN losses by comparing two models: one (PriorDDPM) trained without GAN and the other (PriorDDPM-GAN) trained with GAN (more specifically, the PriorDDPM was trained first till full convergence of its training losses, and then the PriorDDPM-GAN was tuned from the trained PriorDDPM by using extra learning epochs with the GAN losses). Two types of metrics have been considered: one type were the structural deviations between the OSs and the natural ISs, and the other type were the so-called self-consistent structure deviations, which are the deviations between the OSs and the structures predicted from sequences designed on the OSs (here we applied ABACUS-R, a deep learning method for fixed backbone sequence design [16], to select amino acid sequences on the OSs, and used AlphaFold2 to predict the potential folded structures from the selected sequences). The results in Figure 2B shows that PriorDDPM-GAN outperforms PriorDDPM by large margins in both types of metrics. Notably, the low self-consistent RMSD (on average 1.59 Å and the relatively high predicted local distance difference test (pLDDT) scores (on average 82.4) of the predicted structures point towards the high physical plausibility of the OSs from the PriorDDPM-GAN.

Figure 2C compared the residue type distributions of the ABACUS-R-designed sequences on the OSs with those designed on the natural structures. The overall distributions are very similar. In figure 2D, the distributions of the per residue ABACUS-R logits scores of the various designed amino acid sequences are compared. The results show that the sequences designed on a significant portion of OSs from PriorDDPM-GAN are of comparable logits scores to the sequences designed on the natural backbones. For the OSs from PriorDDPM, the same scores are clearly lower (worse) than the scores for the natural backbones. Thus tuning the model with the GAN losses has substantially improved the physical plausibility of the denoised OSs.

### 2.3 In silico evaluation of generated structures

We have mainly used the aforementioned self-consistent structural deviations to evaluate the designability or physical plausibility of the backbone structures generated by our model. Three types of generation tasks have been considered. The first type of task is to use natural backbones as ISs (as has been described in the previous section) and it is mainly for testing purposes. The second type of task is to use a sketch coarsely following a given overall topology as ISs. The OSs of this type of task correspond to de novo backbones, some of which adopt novel, non-natural folded topologies. The third type of task is to inpaint masked regions of a natural backbone. Depending on the size and location of the masked region, this type of tasks may correspond to applications from loop redesign to functional motif scaffolding [19].

Figure 3 summarizes the results of using natural backbones as ISs. For the 25 natural ISs, the median self-consistent RMSD (scRMSD) values is 1.43 Å, while the median self-consistent template alignment-score (scTM-score) is 0.92. The pLDDT scores of the predicted structures strongly correlates with the scTM-scores with the median pLDDT scores being 85.8. As examples, the OSs and the AlphaFold2-predicted structures of the ABACUS-R sequences for the OSs are compared for two test backbones in Figure 3C. Among the two structures shown in Figure 3C, one (1colA00) is all-*α* and the other (1cmsA02) is mainly *β*. The remaining test backbones span other general topology types.

**Figure 3.**
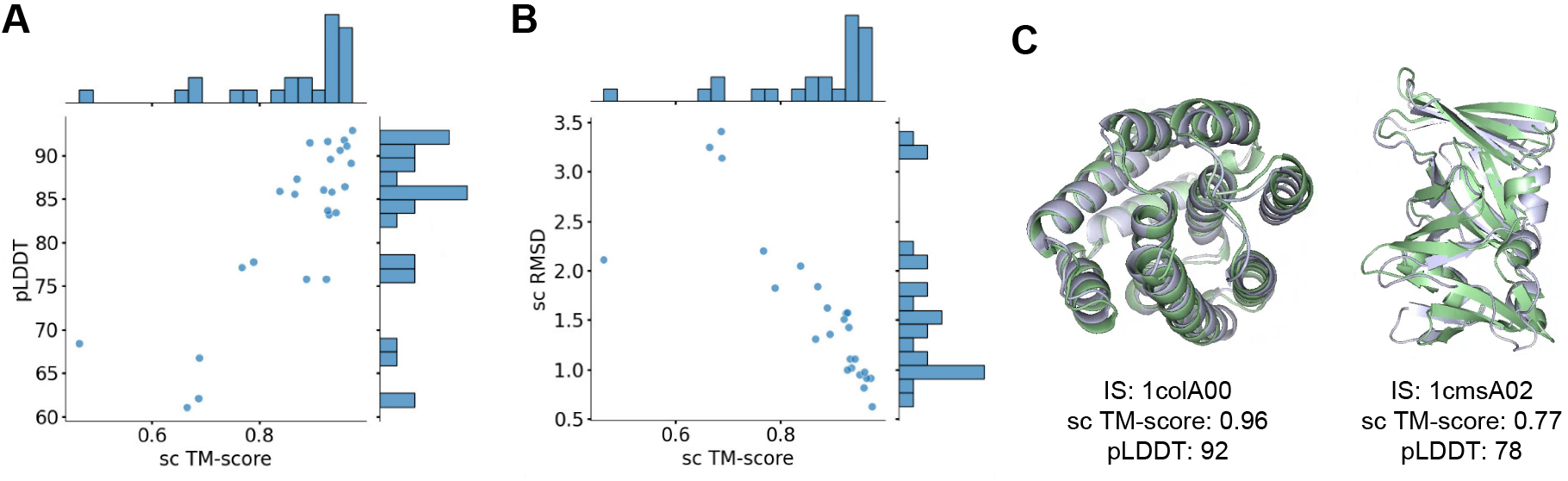
In silico evaluation of output structures produced from natural initial structures. Each natural structure is “denoised” one time to produced a single output structure. The deviations of an output structure from AlphaFold2-predicted structures for a ABACUS-R-designed sequence on that structure are denoted by scTM-score and scRMSD. The pLDDT scores reflect confidence of the AlphaFold2 predictions. **A**, Scattering plot and distributions of the scTM-scores and pLDDT scores. **B**, Scattering plot and distributions of the scTM-scores and scRMSDs. **C**, Two examples of output structures (green) superimposed with AlphaFold2-predicted structures (grey). The CATH protein domain ID of the corresponding initial structure (IS) is indicated under each image.

Figure 4 shows the results of using coarse backbone sketches as ISs. For each of the 125 tested ISs, three Oss were generated using independent denoising runs, and the results for the OS that leads to the highest scTM-score are presented. The median scRMSD, scTM-score and pLDDT score are 2.00, 0.83 and 78.3, respectively. The results indicate that a substantial portion of the de novo backbones should be physically plausible or designable.

**Figure 4.**
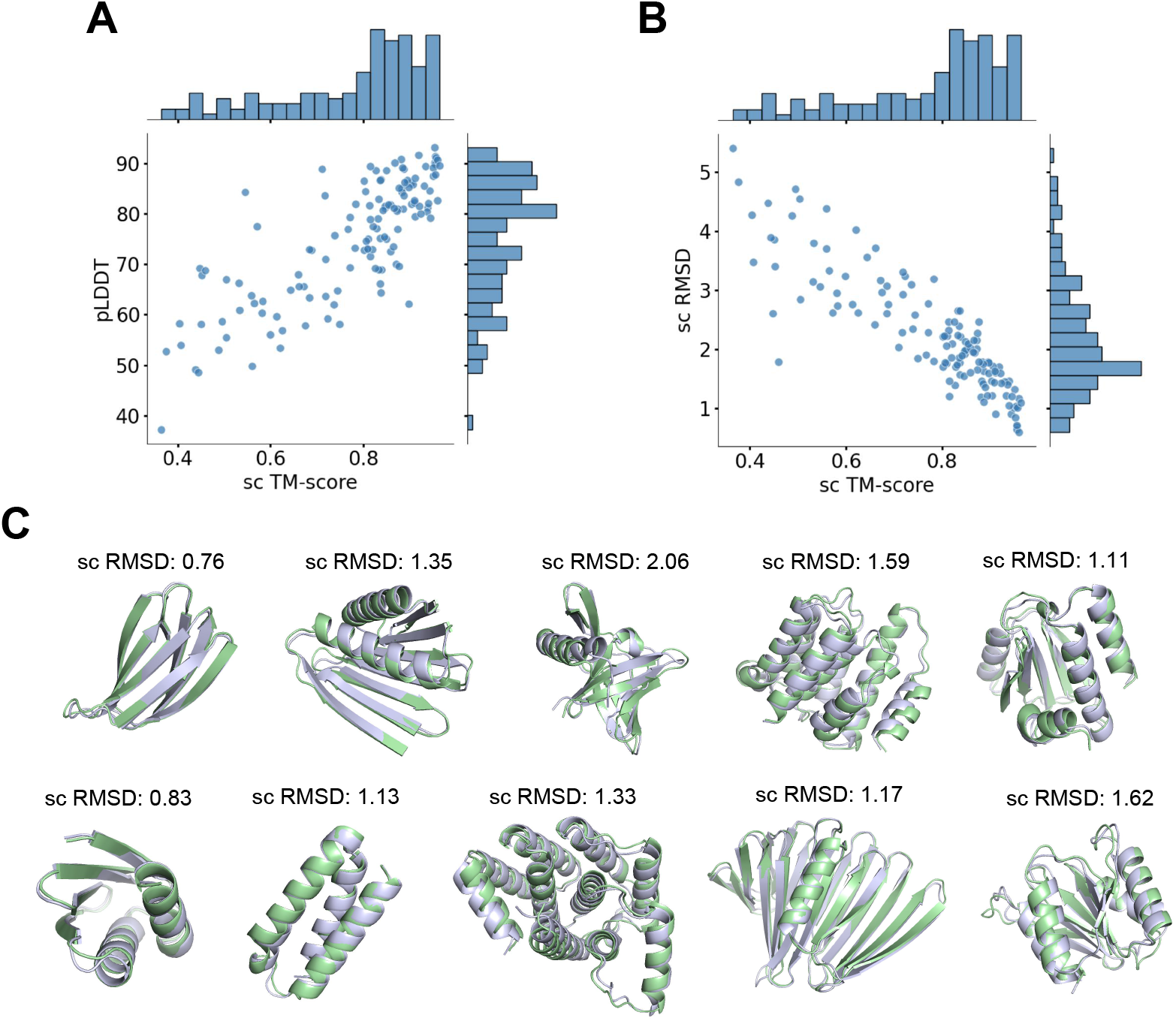
In silico evaluation of output structures having sketched topology. The notations are of the same meaning as in Figure 3. Each initial structure has been denoised three times to generate three output structures, and the results of the largest scTM-score are shown. **A**, Scattering plot and distributions of scTM-scores and pLDDT scores. **B**, Scattering plot and distributions of scTM-scores and scRMSDs. **C**, Examples of output structures (green) superimposed with corresponding AlphaFold2-predicted models (grey). The shown structures are associated with pLDDT ≥ 80 and sc TM-score ≥ 0.8.

Figure 4C shows example backbones generated with the sketched ISs. The OSs are shown together with the AlphaFold2-predicted structures to indicate their potential physical plausibility. Again, these examples illustrate that the current DDPM model with GAN losses can generate a diverse range of backbones.

Figure 5 shows the results of the inpainting tests. We considered six different sizes of the masked window ranging from 5 to 60 residues. For each of the 25 test natural proteins and a given window size, the location of the masked window was randomly selected for three times. For a selected location, we masked the positions of the residues inside the window in the IS and generated three OSs using independent denoising runs. The presented results correspond to the structure of the lowest RMSD from the natural structure among the three OSs. Figure 5A shows that for smaller window sizes (5 or 10 residues), the inpainted structure can well reproduce the natural structure. For larger window sizes, the deviations from the natural structure may no longer be a suitable indicator of the quality (or physical plausibility) of the inpainted structure. Again, we used the self-consistent structural deviations as indicators, which are shown in Figure 5C and 5D for the window size of 60 residues. The median scRMSD, scTM-score and pLDDT score are 1.64, 0.88 and 85.2 As examples, Figure 5B shows several structures from the inpainting tests.

**Figure 5.**
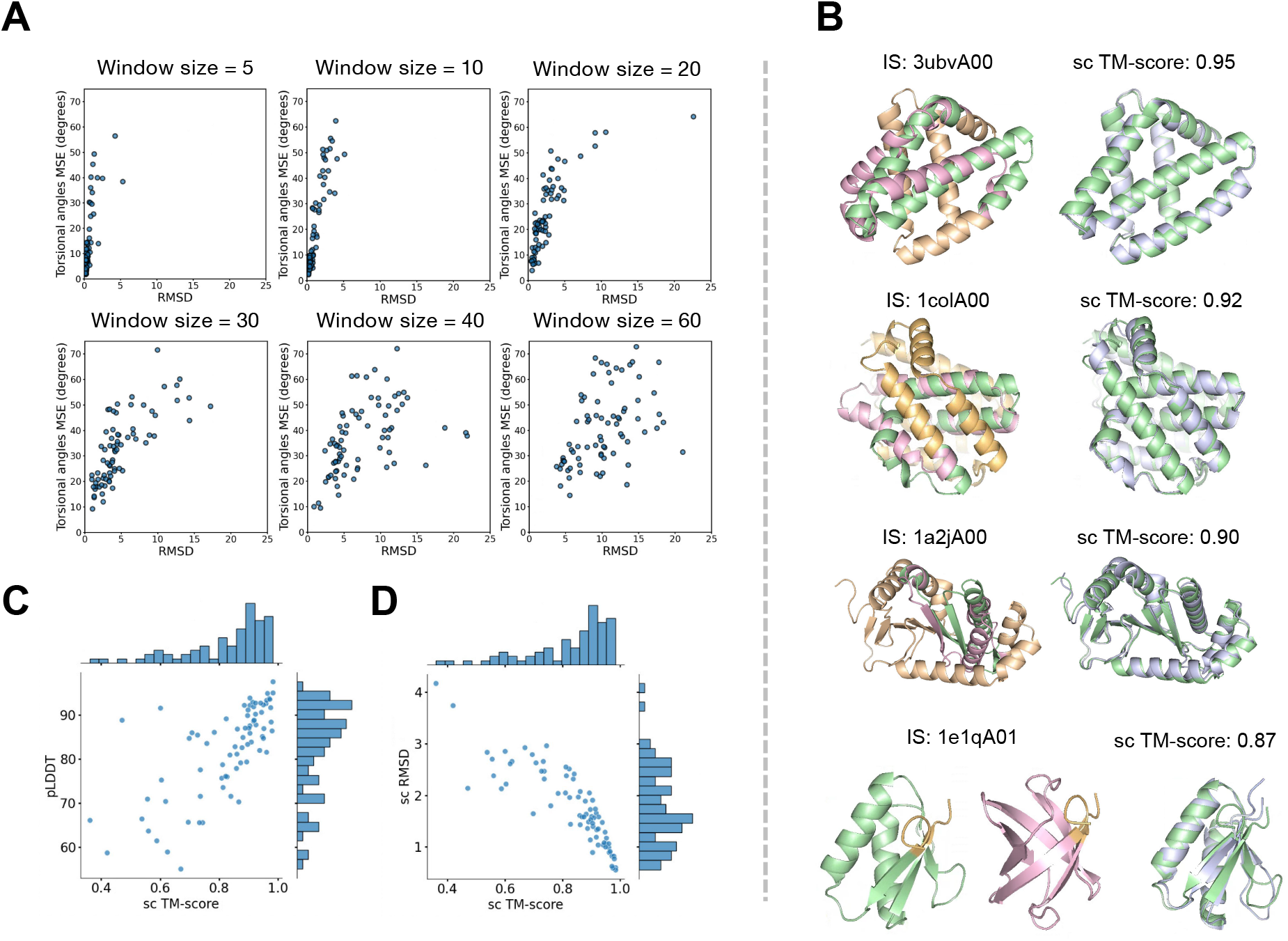
In silico evaluation of the output structures in the inpainting tests. **A**, Deviations of the inpainted structures from the structures of the corresponding regions in the natural structures. Each panel shows the results for a particular size of the masked window. RMSD refers to the deviation of backbone atom positions, tors
ional angle MAE refers to the mean absolute deviation of backbone torsional angles. **B**, Some examples of inpainted structures superimposed with the original structures (left) and with AlphaFold2-predicted structures (right) on ABACUS-R-designed sequences for the inpainted structures. For these examples, the window size of the masked region is 60 residues. The inpainted structures are shown in green, the original natural structures in pink, the preserved (i.e., not masked) regions in orange, and the AplhaFold2-predicted structures in grey. The CATH protein domain IDs of the original structures are given. **C**, Scattering plot and the distributions of scTM-scores and pLDDT scores of the inpainted results for masked window size 60. **D**, Scattering plot and the distributions of scTM-scores and scRMSDs of the inpainted results for masked window size 60. The notations in **C** and **D** are of the same meaning as in Figure 3.

## 3 Discussion

Compared with standard DDPM, SCUBA-D contains several adjustments/extensions to improve the physical plausibility of the denoised structures.

First, the challenging task of generating high quality protein backbones has been divided into two stages. At the firs stage, the partially denoised backbones only need to be coarsely correct or of low resolutions. Thus an efficient single-step LRDM is used for the job. The task of further refining the backbones towards higher resolutions or quality is left to the second stage, in which the SDM takes full advantage of DDPM’s superior ability of synthesizing high quality data. With PriorDDPM, the reverse diffusion process does not need to start from a completely random structure (as in standard DDPM). Instead, it can start from a point that is reasonably close to a plausible backbone. This may significantly reduce the complexity of the problem faced by the SDM, increasing its efficiency in both learning and inference. Moreover, the SDM module can be designed and trained to focus on improving finer structural details which are important for a physically plausible protein backbone. The results in Figure 6A illustrate that the dividing of jobs between the LRDM and the SDM modules works as expected. In the tests of denoising sketched backbones following coarsely defined topologies, the RMSDs between the output structures (OSs) and the initial structures (ISs) were larger than the RMSDs between the prior strctures (PSs) and the ISs, indicating that the LRDM almost always brought the ISs to PSs that are closer to the final OSs than the ISs.

**Figure 6.**
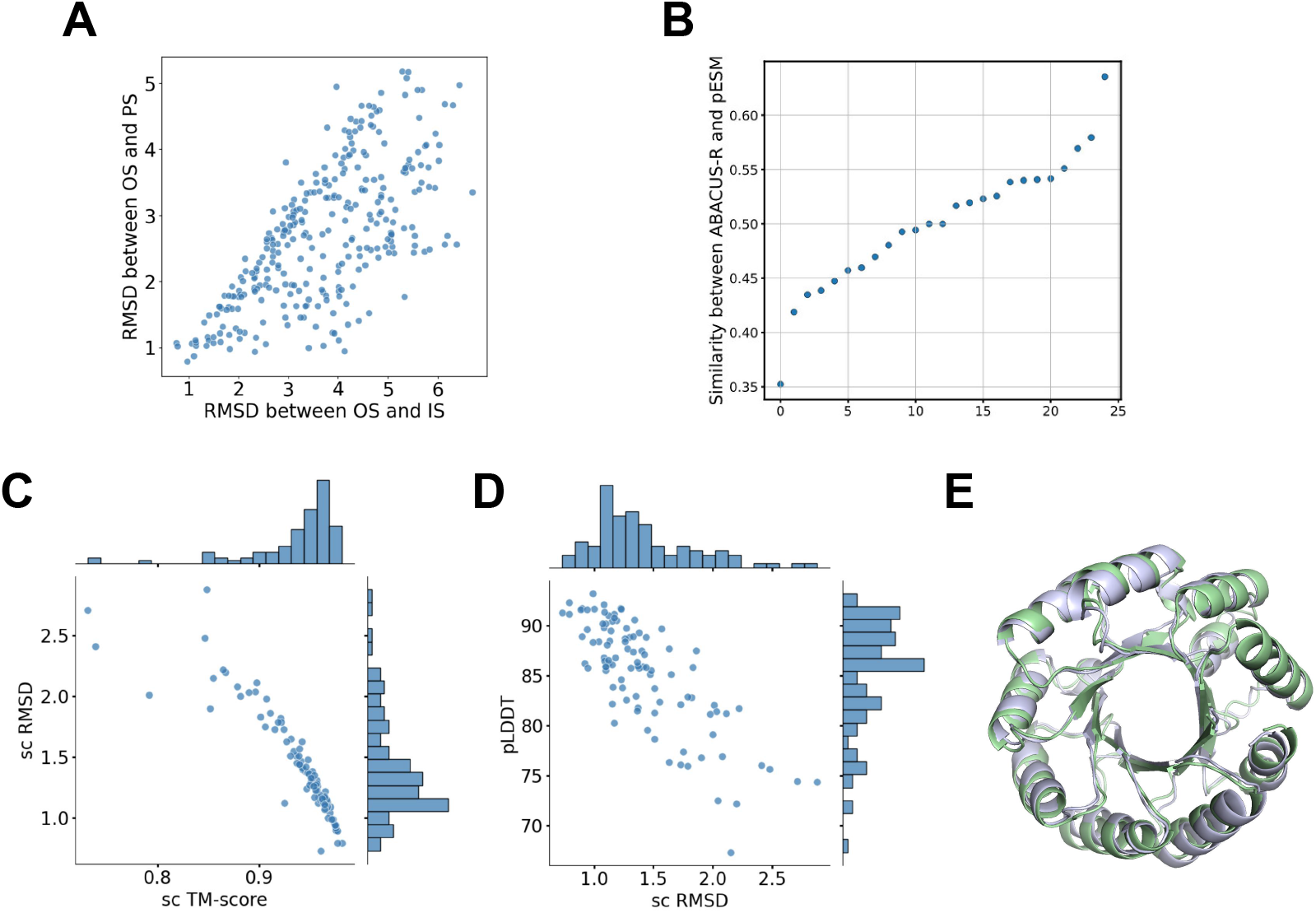
Effects of the various components in SCUBA-D in in silico evaluations. **A**, Scattering plot of RMSD between output structures (OS) and prior structures (PS) and RMSD between output structures (OS) and initial structures (IS). **B**, The similarity between the ABACUS-R-designed sequences and the sequences projected from the output ESM single representation. The projection from the single representation to residue types was carried out using the learned network for determining the cross entropy sequence loss. To compute the sequence similarity, the 20 residues types are divided into 7 groups (GAVLI, FYW, CM, ST, KRH, DENQ and P), with residue types from the same group considered to be similar. Figure 6B shows the sequences from two approaches converged well, which further proves the effectiveness of LM-assisted structure diffusion. The similarity is computed based on 7 coarse residue types. **C**, Scattering plot and the distribution of scTM-scores and scRMSDs. **D**, The same as (C) but for scRMSDs and pLDDT scores. **E**, An example of the output structure (green) and the AlphaFold2-predicted structure. For **A**, the ISs are the various sketched topologies. For **B**, the ISs are the 25 natural protein structures. For **C** to **D**, the initial structures have been sketched to have the topology of an (*αβ*)_9_ barrel, and the notations are of the same meaning as Figure 3.

The use of the PriorDDPM framework also facilitates controllable backbone design. As shown by the examples of the in silico evaluations, different design specifications including the overall topology or pre-defined partial structures (e.g., function motifs) can be effectively implemented through the DDPM prior. This leads to high flexibility of the two-stage PriorDDPM model for handling various types of protein design tasks.

A notable aspect of SCUBA-D is the use of language model (LM)-assisted structure diffusion. Because protein sequences are composed of discrete residue types, they cannot be used directly as the variables in a DDPM, which employs Gaussian noises on continuous variables. Here we replaced the discrete sequences with continuous LM-based single representations. This treatment is similar to Diffusion-LM [39] which embeds discrete tokens into a learnable continuous space and applies denoising diffusion in that space (another way of addressing this problem is the approach of the BERT diffusion model[40], which treats the forward diffusion and the reverse denoising process as a problem of masking and reconstructing tokens similar with the mask language model. A similar idea has also been applied in protein structure and sequence co-diffusion [28]). The results in Figure 2A suggested that incorporating the LM representation definitely improved the recovery of natural structures by the SDM. Intuitively, by requiring the structure diffusion process to reproduce LM-based single representations derived from native sequences, the SDM may better learn how to synthesize a structure with natural protein-like local environments to accommodate different types of side chains, and eventually improve the physical plausibility of the denoised OS. Figure 6B shows that the amino acid sequences derived from the last single representation of SDM are generally similar to the ABACUS-R sequences designed on the OS. This result supports that the single representations and the denoised structures are connected in some physically meaningful ways.

We have used GAN-style losses with two discriminators to further improve the quality of the OSs. This was motivated by the observation that for backbones denoised by the DDPM trained without the GAN losses, their ABACUS-R-designed sequences are associated with lower (worse) logits scores than the sequences designed on the natural backbones (Figure 1D). As the ABACUS-R logits scores reflect the compatibility between side chain types and the local environments surrounding individual residues, the lower scores means distorted local environments in the OSs. We also examined the local backbone conformation distributions of the OSs. Although the lower dimensional marginal distributions (e.g., of one or two torsional angles) could well produce the corresponding distributions of natural backbones, higher dimensional distributions exhibited substantial distorsions (results not shown). Thus considering only structure deviation losses seemed to be unable to provide sufficient restraints on the higher dimensional joint distributions of important structural variables. Our previous study have indicated that to correctly capture the correlations in the joint distributions of different structure features is critically important for sampling protein backbones with high designability[4]. Thus the discriminators for GAN have been specifically engineered to ensure that the model can reproduce the joint distributions that are believed to be important for backbone designability. The results in Figure 1B to 1D indicate that incorporating these GAN losses significantly improved the physical plausibility of the denoised structures.

In summary, we have reported a promising deep learning model for de novo protein backbone design. In in silico evaluations simulating a variety of protein design tasksthe model consistently generated high quality de novo backbones (with high self-consistent TM-scores and pLDDT scores after sequence design). We note that our model remains to be validated by wet experiments.

## 4 Method

### 4.1 The basic denoising diffusion probabilistic model

In traditional areas of machine learning, various deep neural network-based models for generating high quality samples have been proposed, including normalizing flows [41], variational auto-encoders (VAE) [22], generative adversarial networks (GAN) [24] and denoising diffusion probabilistic models (DDPM) [27, 42, 43], etc. Our model is based on DDPMs, which have achieved great successes in generating high quality and semantically editable images[30].

In DDPMs, the process of adding noises to data (referred as the forward process) is defined as a Markovian diffusion process, while the denoising process is modeled with deep networks optimized to reproduce the conditional distributions in the reverse Markovian process.

In the forward process, the initial distribution *q*(***x***_0_) of the observed data ***x***_0_ is gradually converted into a pre-defined distribution (usually a standard normal distribution) by adding noises through a Markov chain of *T* discrete steps. The transition probability from step *t*−1 to step *t* (*t* ∈ [1, 2, …, *T*]) is usually defined to be a normal distribution, namely, 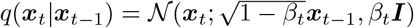, in which *{β*_*t*_, *t* ∈ [1, 2, …, *T*]*}* is called the noise schedule.

The reverse (denoising) process generates a new data sample ***x***_0_ from a noise sample ***x***_*T*_ ∼ *p*(**x**_*T*_) = 𝒩(**0, *I***) with the learned transition probability 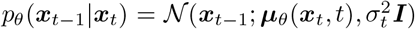, where *θ* is the model parameter and 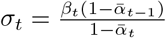 with *α*_*t*_ = 1 − *β*_*t*_ and 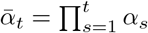.

The standard DDPMs are trained by maximizing the evidence lower bound (ELBO) of the log likelihood of the training data, with the ELBO being the sum of the negative Kullback-Leibler(KL) divergences between the forward and reverse transition distributions of all the diffusion steps[27],

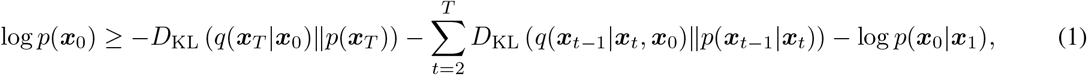

It can be shown that maximizing the above ELBO is equivalent to minimizing the following denoising loss [44],

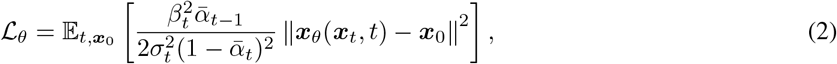

where ***x***_*θ*_(***x***_*t*_, *t*) is the predicted denoised data (noted as 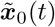 below) based on ***x***_*t*_,, During training, the noise sample ***x***_*t*_ is generated by forward diffusion from a training sample ***x***_0_:

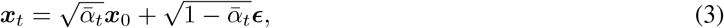

with ***ϵ*** ∼ 𝒩(**0, *I***).

The denoising version of the DDPM loss simplifies the learning process into learning a denoising model ***x***_*θ*_(***x***_*t*_, *t*) that can generate the predicted data 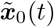 to best reproduce the true data ***x***_0_.

With a denoising model trained with the above scheme, new samples or synthesized data can be generated by iteratively apply the following two steps, starting from sampling the random distribution *p*(***x***_*T*_):

1. predict the denoised sample for step *t* as 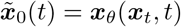;
2. predict the diffused sample at step *t* − 1 by adding noises to the denoised sample for step *t*, namely, 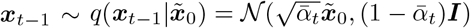.

We note that when ***x***_0_ corresponds to 3-D structures, the denoised structures may be superimposed onto the same reference structure before adding noises in step 2 to remove potential overall translation and rotation. This, however, should not be necessary when the network ***x***_*θ*_(***x***_*t*_, *t*) is SE(3) equivariant[28].

In this study, we adopted a recently proposed variant of the above standard DDPM model, the PriorDDPM [32, 33], for generating protein backbone structures. In PriorDDPM, the end point of the forward diffusion process is defined to follow a Gaussian distribution with non-zero means, i.e. ***x***_*T*_ ∼ 𝒩(***μ, I***). This is achieved by modifying equation (3) into

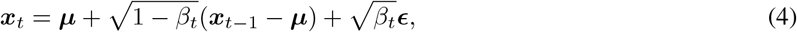

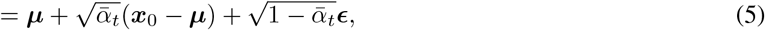

Similar to standard DDPM, the PriorDDPM can be trained with a denoising loss as defined in (11). During training and inference, the denoising network can now be guided by the prior, namely, 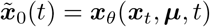.

With PriorDDPM, the diffusion process is not required to connect real data with noised data centered around the origin. Instead, it only needs to connect real data with noised data centered around a given prior (i.e., ***μ***). If we assume that the priors will always be reasonably close to real data, both the training of and the inference with PriorDDPM can be much more efficient than standard DDPM.

We have used a separate network that works like a denoising auto-encoder to generate the prior backbone structure in a single step. As the prior is to be further refined by the downstream DDPM, this auto-encoder only needs to produce a low resolution backbone, which corresponds to ***μ*** in Equation 5.

### 4.2 Modelling protein backbone structures

We describe a protein backbone structure as a collection of the positions and orientations of individual residues. For each residue, its position and orientation can be specified by a rigid body transformation 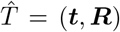, in which the vector ***t*** ∈ ℝ^3^ denotes translation of the C_*α*_ atom coordinates relative to the origin of a global coordinate frame, while ***R*** ∈ SO(3) denotes the rotation of a local coordinate frame (i.e., a coordinate frame fixed on the residue) relative to a fixed reference frame in the global coordinate system. The rotation can be specified with a quaternion ***q*** = [*q*_1_, *q*_2_, *q*_3_, *q*_4_], which can eventually be considered as a feature vector of 3 angular values, namely,

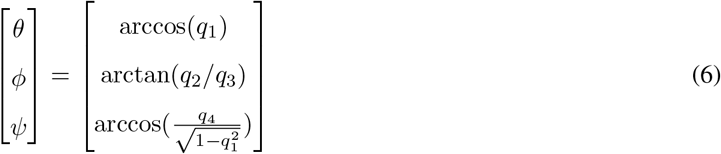

The structure diffusion is performed in the space spanned by the above feature vectors, with the intra-residue bond lengths and bond angles assumed to be fixed.

Following the ideas of AlphaFold2 of transforming protein structures with SE(3) equivariant deep networks, we first transformed the above residue-wise (***t, R***) features (describing an input structure containing noises) into a residue pair-wise features, which correspond to the relative translations and rotations between each pair of residues[20]. We used triangle attentions to update the pair representation followed by Invariant Point Attention[20] to retrieve the residue-wise (***t, R***) representation of the (denoised) output structure. This architecture has been adopted to implement the structure denoising networks in both the LRDM and the SDM.

We have used sequence representations to assist the structure diffusion in SDM. For this purpose, we used the ESM represenation, which is a language model pre-trained from massive samples of known natural protein sequences [35]. The LRDM comprises an extra decoder which predict an ESM-based representation (the single representations shown in Figure 1B) of the amino acid sequence from the denoised prior structure (PS). This decoder has been implemented as a GVP subnetwork. The output single representation is used as the prior for the single representation in the downstream DDPM module (the SDM). In SDM, the ESM-based single representation is refined in the same way as the structure, namely,

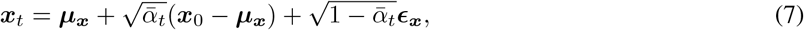

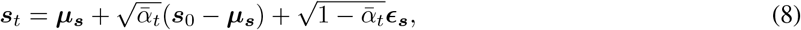

in which ***μ***_***x***_ and ***μ***_***s***_ denote respectively the priors of the structure and of the sequence representation. The denoising subnetworks can be noted as

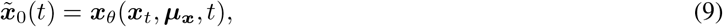

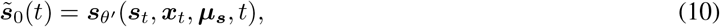

in which the subnetwork updating the single representation receives input from both the pair and the single representation as in AlphaFold2. However, when the pair representation is updated, directly using the information of the single representation as in AlphaFold2 was not found to be beneficial, probably because the single representation here has only been predicted from a low resolution prior structure instead of being derived from an actual amino acid sequence or sequence profile. Thus the single representation was not directly involved in Equation 9

During training, the contributions of the LRDM to the total loss include the frame align point error or FAPE loss (FAPE_prior_) computed using the output of the structure denoising subnetwork and the mean squared error loss (MSE_prior_)computed on the output ESM-based single representation. For the mutimodal DDPM module (the SDM), the ELBO-based denoising loss for step *t* should be defined as

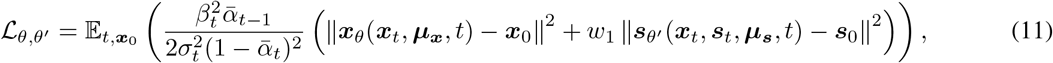

In our final model, we have made the following adjustments to this total loss. Firstly, we have replaced the MSE-based structural loss ∥***x***_*θ*_(***x***_*t*_, ***μ***_***x***_, *t*) − ***x***_0_∥^2^ by structure deviation losses supplemented with GAN-style losses (see below). The structure deviations losses comprised the FAPE loss (FAPE_DDPM_), a distogram classification loss (CE_dist_), and a covalent geometry violation loss (loss_violation_). The CE_dist_ was computed by predicting (using a learned subnetwork) the C_*α*_ distance map from the last layer pair-representation of the denoising subnetwork, and comparing the predicted map with the actual map (to compute the loss as the sum of classification errors over all distances, the distance range within 20 Å was evenly divided into 64 bins and each bin was assigned a distance class). The loss_violation_ is the mean squared deviations of bond lengths and bond angles from chemically allowed ranges. Secondly, we have supplemented the MSE-based sequence representation loss (MSE_DDPM_) with an extra cross entropy loss (CE_aatype_) between the actual amino-acid sequence and an amino acid sequence predicted from the denoised ESM-based representation. The latter extra loss has been included to ensure that the output single representations come from a semantically reasonable subspace, and can be noted as *w*_2_ Cross Entropy(NN_sequence_(***s***_*θ*′_), sequence_0_), in which *w*_2_ is an empirical weight, and NN_sequence_(·) denotes the sequence prediction network, which is a learned projection head that projects the single ESM representation at each position to 20 residue types.

### 4.3 The discriminator networks for GAN-style losses

We used two discriminator networks to provide GAN [24] losses to supplement the structure deviation losses in the SDM module. One is a graph-based packing discriminator, the other is a transformer-based backbone torsion correlation discriminator. These two discriminators evaluate the quality or physical plausibility of the denoised structure from two aspects: the first discriminator mainly evaluate the local packing around individual residues, while the second discriminator mainly evaluate the backbone conformations of contiguous peptide segments.

For the graph-based packing discriminator, the backbone structure is modeled as a graph, with nodes corresponding to amino acid residues and edges connecting spatially neighboring residues. The node features include the local backbone conformation represented by the 21 backbone torsional angles from *ϕ*_*i*−3_, *ψ*_*i*−3_, and *ω*_*i*−3_ to *ϕ*_*i*+3_, *ψ*_*i*+3_, and *ω*_*i*+3_, which are transformed with the sin and cos functions. The edge features include distances between the N, C_*α*_, C, and a virtual C_*β*_ atoms of the two residues (these are similar to the edge features used by the sequence design program ProteinMPNN [17]). Besides these, the edge features also include a 7-component vector ***O***_*ij*_ = [***t***_*ij*_, ***q***_*ij*_] describing relative orientations between the local coordinate frames constructed from three consecutive C_*α*_ atoms (i.e., 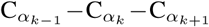, with *k* ∈ *{i, j}*). More specifically, ***t***_*ij*_ is the normalized translation vector and ***q***_*ij*_ the rotational quaternion from the local frame at *i* to the local frame at *j*, both represented in the local frame at *i*.

The backbone torsion correlation discriminator is a transformer-based model, which encodes the backbone torsion angles (*ϕ, ψ*) sequentially along the peptide chain by using global attention.

The discriminator losses of the GAN are defined as

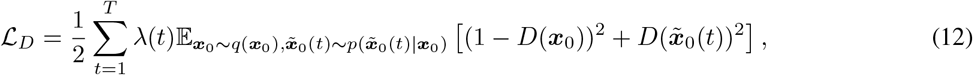

where D(·) denotes the discriminator, and 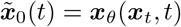 is the generator in the context of GAN modelling.

The generator losses (loss_gen_) are defined as

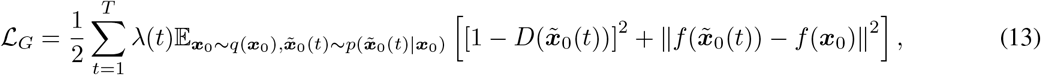

where f(·) denotes the values of the internal neuron nodes of the discriminator. Following the DDPM framework, the GAN losses are weighted by 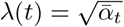.

### 4.4 Training with natural protein structures

Our model has been trained by learning to denoise perturbed natural protein structures. To let the model learn how to recognize a diverse range of initial structures that the model will potentially run into in various downstream design tasks, we used the following 4 different strategies to obtain initial structures by perturbing the training natural structures.

S1: masking the structure information of a certain fraction of residues from a natural structure. The fraction of masked residues was random selected to be between 10% to 30% in the initial phase of training and increased to between 10% to 70% in the fine tuning phase (see blow) of training. The masked residues can be either multiple segments of 3 to 7 residues (substrategy S1-1), a single long segments (substrategy S1-2), or a set of residues that are spatial neighbors surrounding a randomly selected residue (substrategy S1-3). In the ISs, the masked residues take random positions (from a normal distribution with zero mean and 15 Å variance) and orientation (uniform distribution)).

S2: adding random noise to the location and orientation of every residue in a natural structure.The noise vector added to the residue location was sampled from a normal distribution with zero mean and a variance of 3 Å. The orientation was sampled from the uniform distribution.

S3: adding random noises to the location and orientation of every secondary structure element in a natural structure. An entire secondary structure element is translated by a random vector sampled from a normal distribution with zero mean and a variance of 3 Å. The orientation is changed by interpolating between the natural orientation (***v***_0_) and a random orientation (***v***_1_) sampled from the uniform distribution, namely:

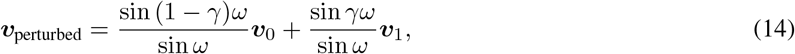

where *ω* is the angle between the rotational axes, and the parameter *γ* takes the value of 0.5 here. With the secondary structure elements perturbed, all residues in loops are treated as being masked.

S4: generating perturbed backbones by running stochastic dynamics (SD) simulations with SCUBA starting from a natural structure. The potential energy function used for the SD included the covalent and Van der Waals energy terms but no other energy terms of SCUBA. The temperature of the simulations was set to 5 (in *k*_*B*_*T* unit) to encourage the sampling of diverse structures. For each training natural structure, the SD simulations were pre-performed for 100 *ps* and 20 structures evenly spaced in time were extracted from the last 70 *ps* of the simulation.

### 4.5 Initial structures sketched according to given topologies

To evaluate the performance of model for refinement from coarse backbone sketches, low resolution structures following given topologies were built with a coarse grain model for positioning secondary structure elements (Miao et al, unpublished) and the SCUBA backbone sketch tool[4]. The compositions of secondary structure elements in these ISs were the same as the 25 natural domains used in the natural IS tests.

The ISs for the (*αβ*)_9_ designs were built first by using the SCUBA backbone sketch tool to arrange 9 peptide segments in the helical conformation and another 9 segments in the strand conformation in a double ring configuration Loops were subsequently added to connect the segments. We found that if these sketches were used as ISs, the chances of obtaining OSs as closed barrels were small. Thus we used restrained SCUBA SD simulations to relax the initial sketched structures. In total 33 SCUBA relaxed backbones were used as the ISs.

### 4.6 More details about the implementation and training

Our model was trained using the structures of natural protein domains defined in the CATH database. The available set of non-redundant natural domains were split into training, validation and test sets with the ratios of 90:5:5. To maintain independence across the sets, any pair of domains from two different sets do not belong to the same topology category (according to the CATH 4.2 classification of protein structures).

At each training step, a part of maximally 128 residues was cropped from a natural structure domain using the same approach as AlphaFold2 (the probabilities of using sequence-based cropping or using spatial location-based cropping is 0.5:0.5). An initial structure was generated by perturbing the cropped structure with one of the strategies among S1-S4 (with approximate probabilities 0.5:0.17:0.17:0.17). If S1 was chosen, the probabilities for choosing the substrategies S1-1 to S1-3 were approximately 0.33:0.33:0.33. The initial structures for strategies S1 to S3 were generated on the fly. If S4 was selected, one initial structure is randomly selected from the stored 20 structures produced by the pre-performed SD simulation.

The training was divided into two phases: an initial phase and a fine tuning phase. The initial phase lasted for 130k training steps (one step corresponds to the denoising of one initial structure). The GAN losses were not considered in this phase. The fine tuning phase was started with the parameter values learned in the initial phase, and the GAN losses were considered in this phase, which also lasted 130k steps. The entire training took around 3 weeks on 24 V100 12G GPUs. More training details are summarized in Figure 7.

**Figure 7.**
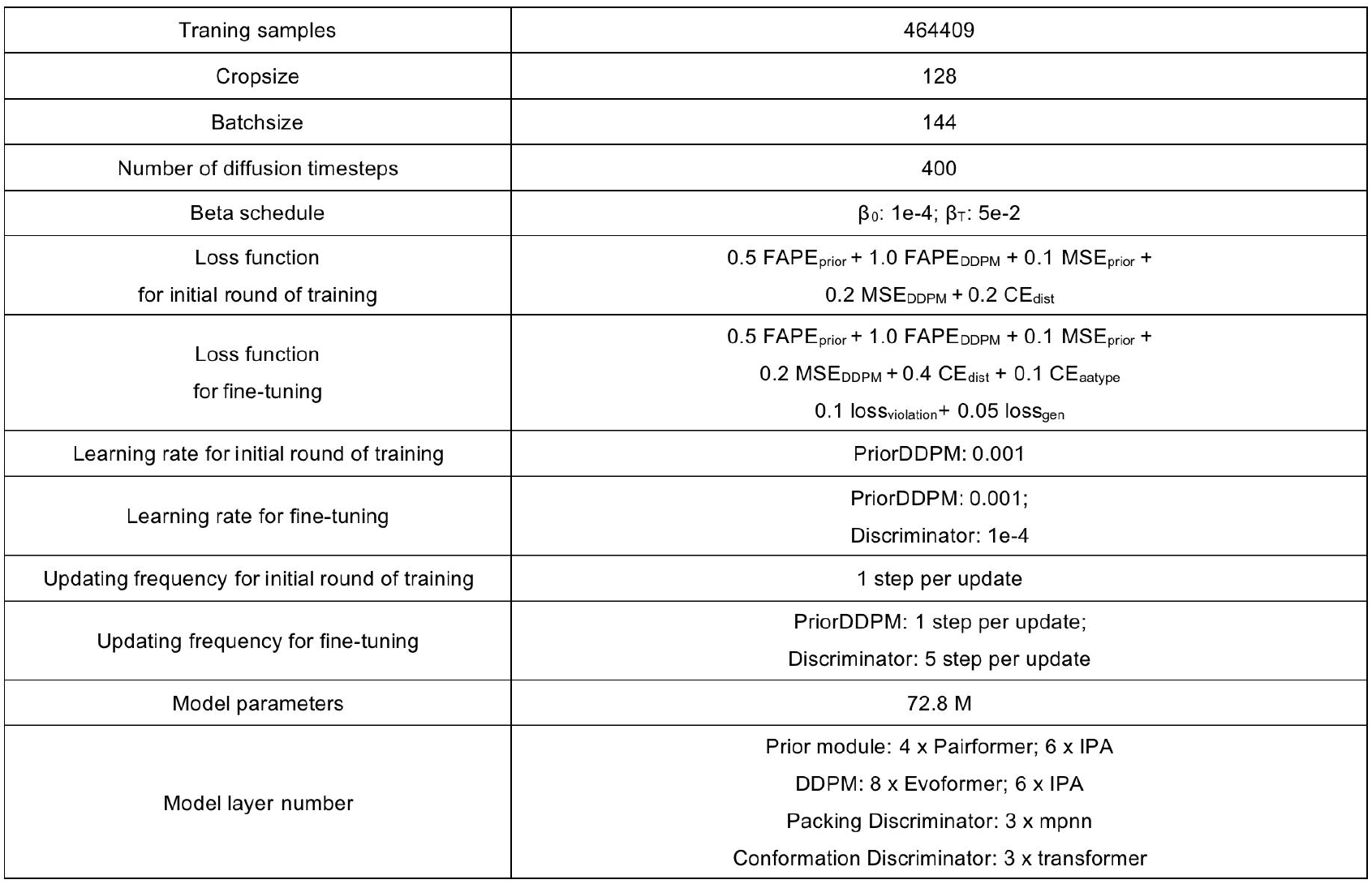
Details of networks and training.

